# Platelets Outperform Leukocytes in Transcriptomic Liquid Biopsy Profiling of Myeloproliferative Neoplasms

**DOI:** 10.64898/2026.03.30.714941

**Authors:** Zhu Shen, Abhishek Sawalkar, Jason Wu, Vanita Natu, Jesse Rowley, Matthew Rondina, Anandi Krishnan

## Abstract

Myeloproliferative neoplasms (MPNs) are characterized by progressive myelofibrosis that drives morbidity and mortality. Liquid biopsy approaches to noninvasively monitor fibrotic progression remain limited. We performed comparative transcriptomic profiling of CD45-depleted platelet-enriched and CD45+ leukocyte-enriched fractions from matched peripheral blood samples of 76 individuals (27 primary myelofibrosis, 17 polycythemia vera, 14 essential thrombocythemia, 18 healthy controls). Platelet RNA sequencing was performed in 2018-2020 on Illumina HiSeq 4000, while WBC RNA sequencing was conducted in 2023 on Illumina NovaSeq 6000 from cryopreserved CD45+ enriched fractions of specimens obtained at the identical time and from the same blood sample as the platelet RNA. Despite comparable library preparation protocols and higher sequencing depth in WBC samples, platelet transcriptomes exhibited 5.1-fold more differential expression in myelofibrosis (3,453 versus 681 genes, adjusted p<0.05, |log2FC|>1). Platelet signatures were enriched for proteostasis pathways including endoplasmic reticulum stress and unfolded protein response, reflecting megakaryocyte dysfunction in the fibrotic bone marrow niche. WBC signatures predominantly featured immune activation and proliferative pathways, indicating systemic inflammatory responses. Multinomial LASSO classification demonstrated superior performance of platelet-based models for myelofibrosis diagnosis (AUROC 0.85) compared to WBC-based (AUROC 0.77) or clinical models (AUROC 0.59). Combined platelet+WBC models did not improve performance (AUROC 0.80), indicating complementary but non-additive information. These findings establish platelet transcriptomic profiling as a superior noninvasive biomarker platform for monitoring myelofibrosis in MPNs, capturing megakaryocyte-driven fibrogenesis with greater sensitivity than peripheral leukocyte-based approaches.

**Highlights:** Using matched WBC and platelet RNA-seq from MPN patients, we identify myelofibrosis-associated transcriptomic signatures specifically enriched in platelets.

Multinomial LASSO modeling highlights platelet-derived gene expression as a dominant and predictive biomarker of myelofibrosis, outperforming clinical parameters and WBC signatures.

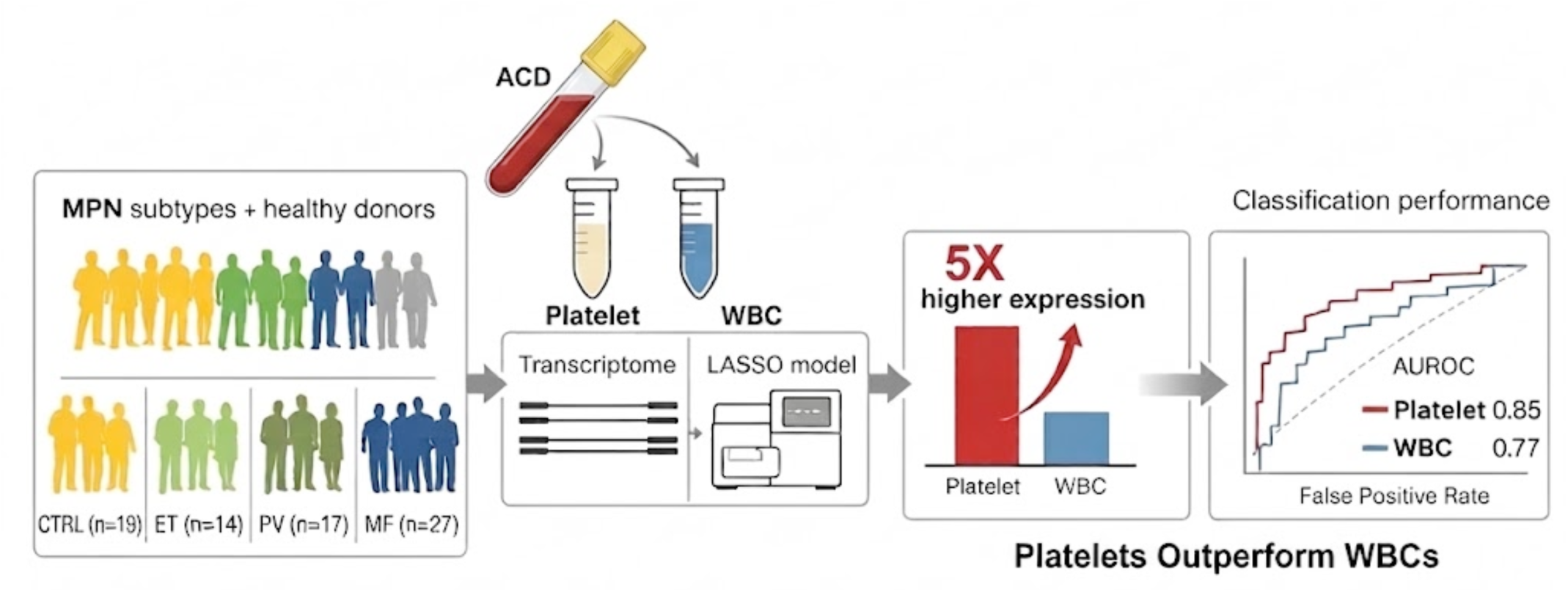

## Introduction

Myeloproliferative neoplasms (MPNs) are a heterogeneous group of Philadelphia chromosome-negative hematologic malignancies characterized by clonal proliferation of myeloid lineage cells and variable degrees of bone marrow fibrosis (myelofibrosis, MF), which significantly contributes to morbidity and mortality^1–5^. The molecular pathogenesis of MF involves complex interactions between driver mutations (e.g., JAK2, CALR, MPL) and the bone marrow microenvironment, including proinflammatory signaling, aberrant cytokine production, and altered stromal cell function^6–13^. Recent advances in transcriptomic profiling, particularly bulk and single-cell RNA sequencing (RNA-seq), have provided valuable insight into cellular and molecular mechanisms underlying MPN disease progression, with megakaryocytes and platelets emerging as key contributors to fibrotic remodeling and disease severity^14–25^.

Platelets, anucleate cytoplasmic fragments derived from megakaryocytes, retain a dynamic transcriptome reflective of megakaryocyte biology and host responses to systemic disease onset and progression. Therefore, platelets are attractive liquid biopsy candidates for noninvasive biomarker discovery in MPNs^26–31^. Several studies have identified platelet-specific gene expression signatures enriched in pathways related to proteostasis, endoplasmic reticulum (ER) stress, and the unfolded protein response; processes implicated in fibrogenesis^32^. In parallel, peripheral white blood cells (WBCs), including CD45+ leukocytes, contribute to inflammation and immune dysregulation in MPNs, but their relationship to fibrosis-associated transcriptional programs remains less defined^33–35^.

Despite these advances, an integrated comparative analysis of matched platelet and WBC transcriptomes within the same MPN patient samples has not been fully explored. Understanding how distinct hematopoietic compartments reflect fibrosis biology may reveal complementary or dominant biomarkers for disease monitoring and therapeutic targeting. Here, we performed transcriptomic profiling of platelet and WBC fractions derived from the same MPN patients, identifying fibrosis-associated, compartment-specific signatures and demonstrating the superior predictive capability of platelet transcriptomes in assessing myelofibrosis severity.

## RESULTS

### Patient Cohort Characteristics and Technical Quality

We performed comparative transcriptomic profiling of matched CD45-depleted platelet-enriched and CD45+ leukocyte-enriched fractions isolated from the same donor blood draws in 76 individuals, including 14 essential thrombocythemia (ET), 17 polycythemia vera (PV), 27 primary myelofibrosis (MF) patients, and 18 healthy controls. Both platelet and WBC fractions were purified from identical blood samples collected between 2017 and 2019, with platelet fractions sequenced immediately (2018-2020, HiSeq 4000) and WBC fractions cryopreserved for later sequencing (2023, NovaSeq 6000). Despite this temporal separation of sequencing runs, both datasets achieved high technical quality with comparable library preparation protocols and read configurations (2×75 bp paired-end, ∼40 million read target). WBC samples detected more genes (16,383 vs 12,911 genes with >10 counts), likely reflecting both the greater complexity of CD45+ leukocyte transcriptomes and the improved sensitivity of a more recent sequencing platform. Consistent with higher transcriptome complexity in leukocytes, sequencing reads in WBCs were distributed across a broader set of genes: the top 5, 10, 20, 100, 500, and 1000 genes accounted for 9.87%, 15.89%, 23.02%, 40.87%, 62.31%, and 72.68% of total WBC reads, respectively, compared with 15.75%, 24.31%, 34.85%, 54.84%, 75.37%, and 83.69% in platelets. Technical variability was comparable between datasets, with median coefficient of variation of 0.72 in platelets and 0.78 in WBCs, indicating similar measurement precision.

The patient cohort reflected real-world clinical diversity, including treatment-naive individuals and patients receiving cytoreductive therapies, JAK inhibitors, or anti-thrombotic agents. Patient age, sex, mutation status, and treatment regimen were balanced across disease subtypes and were adjusted as covariates in all differential expression analyses to isolate disease-specific transcriptional signals from treatment-related effects.

### Platelet Transcriptomes Exhibit Substantially Greater Differential Expression Than WBC Transcriptomes

Independent differential expression analysis revealed markedly greater transcriptional dysregulation in the platelet compartment compared to CD45+ leukocytes across all MPN subtypes (Figure 1). In primary myelofibrosis patients, 3,453 genes were significantly differentially expressed in platelets versus controls (adjusted p<0.05, |log2 fold change|>1), compared to 681 genes in the WBC fraction: a 5.1-fold difference. This pattern was consistent across MPN subtypes: ET patients exhibited 941 differentially expressed genes in platelets versus 549 in WBCs (1.7-fold), while PV patients showed 1,714 versus 557 genes (3.1-fold). The magnitude of compartment-specific divergence was most pronounced in MF, the disease stage characterized by advanced bone marrow fibrosis, suggesting that platelet transcriptomes increasingly capture disease-specific pathology as fibrosis progresses.

**FIGURE 1.**
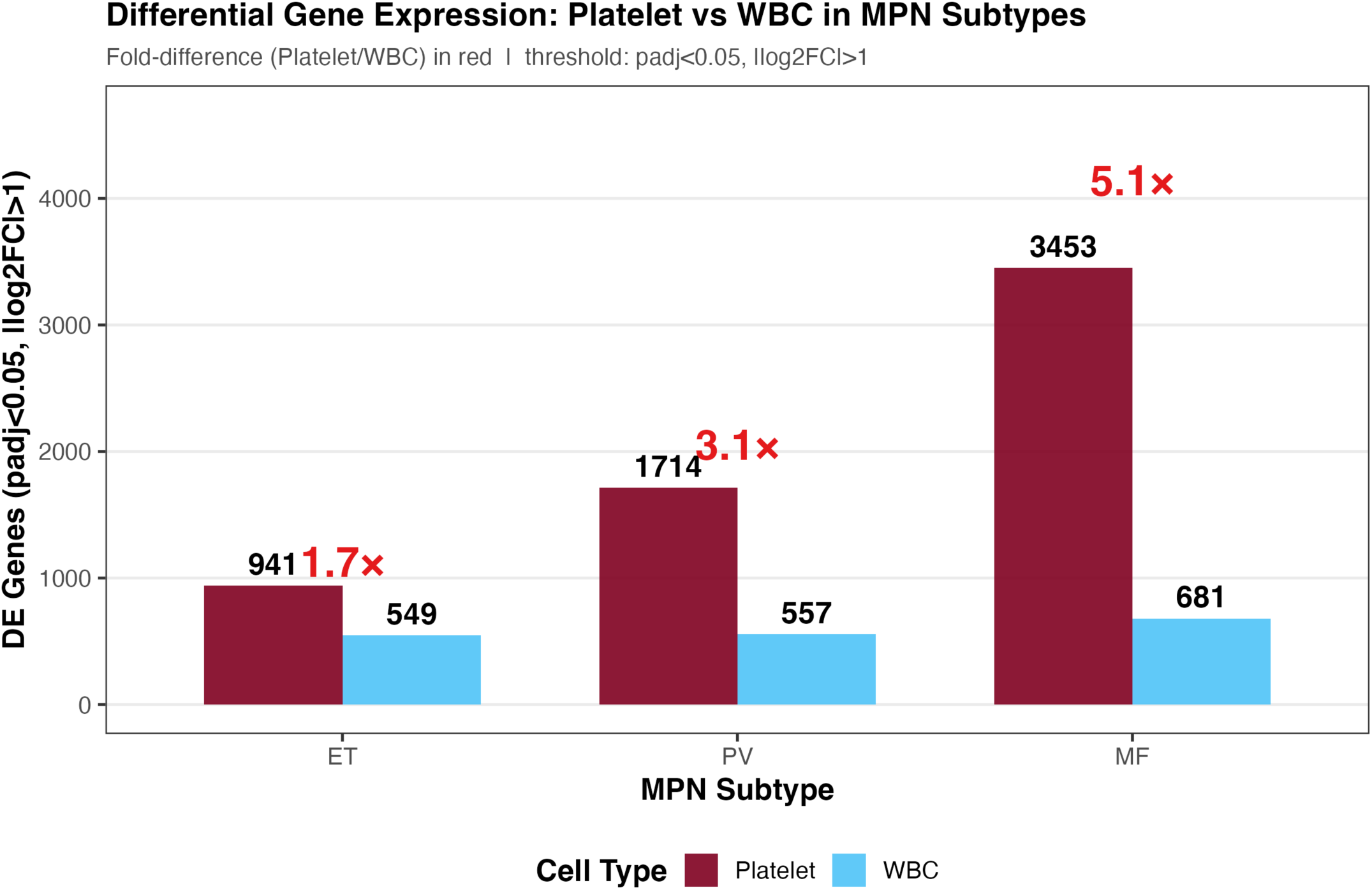
Platelet transcriptomes exhibit greater differential expression than WBC transcriptomes across MPN subtypes. Bar chart comparing the number of significantly differentially expressed genes (adjusted p<0.05, |log2 fold change|>1) between platelet (dark red) and WBC (blue) transcriptomes in essential thrombocythemia (ET), polycythemia vera (PV), and primary myelofibrosis (MF) patients versus healthy controls. Each bar represents the total number of genes meeting differential expression criteria in independent DESeq2 analyses. Numbers above bars indicate gene counts. Red fold-difference values above each disease category show the ratio of platelet to WBC differentially expressed genes. MF demonstrates the greatest divergence (5.1-fold), with 3,453 platelet genes versus 681 WBC genes significantly dysregulated, indicating that platelet transcriptomes capture more extensive disease-related transcriptional changes than WBC transcriptomes, particularly in advanced fibrotic disease.

Direct comparison of log2 fold changes between compartments revealed largely non-overlapping differential expression patterns, with 88.6% of platelet DE genes showing platelet-specific dysregulation (**Supplementary Figure 1, Supplementary Table 1**). This platelet-dominant pattern held across all pairwise disease comparisons, with platelet:WBC DE gene ratios ranging from 1.7-fold to 15.1-fold. The 395 MF genes significantly dysregulated in both compartments exhibited concordant directionality, indicating biological coherence rather than technical artifacts.

Hierarchical clustering using the top 1% most variable genes (129 platelet genes, 164 WBC genes) demonstrated clear separation of disease from healthy controls in both compartments, with additional clustering by MPN subtype most evident in platelets (**Figure 2**). Platelet samples exhibited tight clustering by disease status, with all MPN subtypes (ET, PV, MF) forming distinct transcriptional blocks separate from healthy donors. In contrast, WBC samples showed more dispersed clustering patterns, consistent with greater cellular heterogeneity in the CD45+ leukocyte population. Notably, the gene sets driving disease classification were largely non-overlapping between compartments, indicating distinct biological signatures.

**FIGURE 2.**
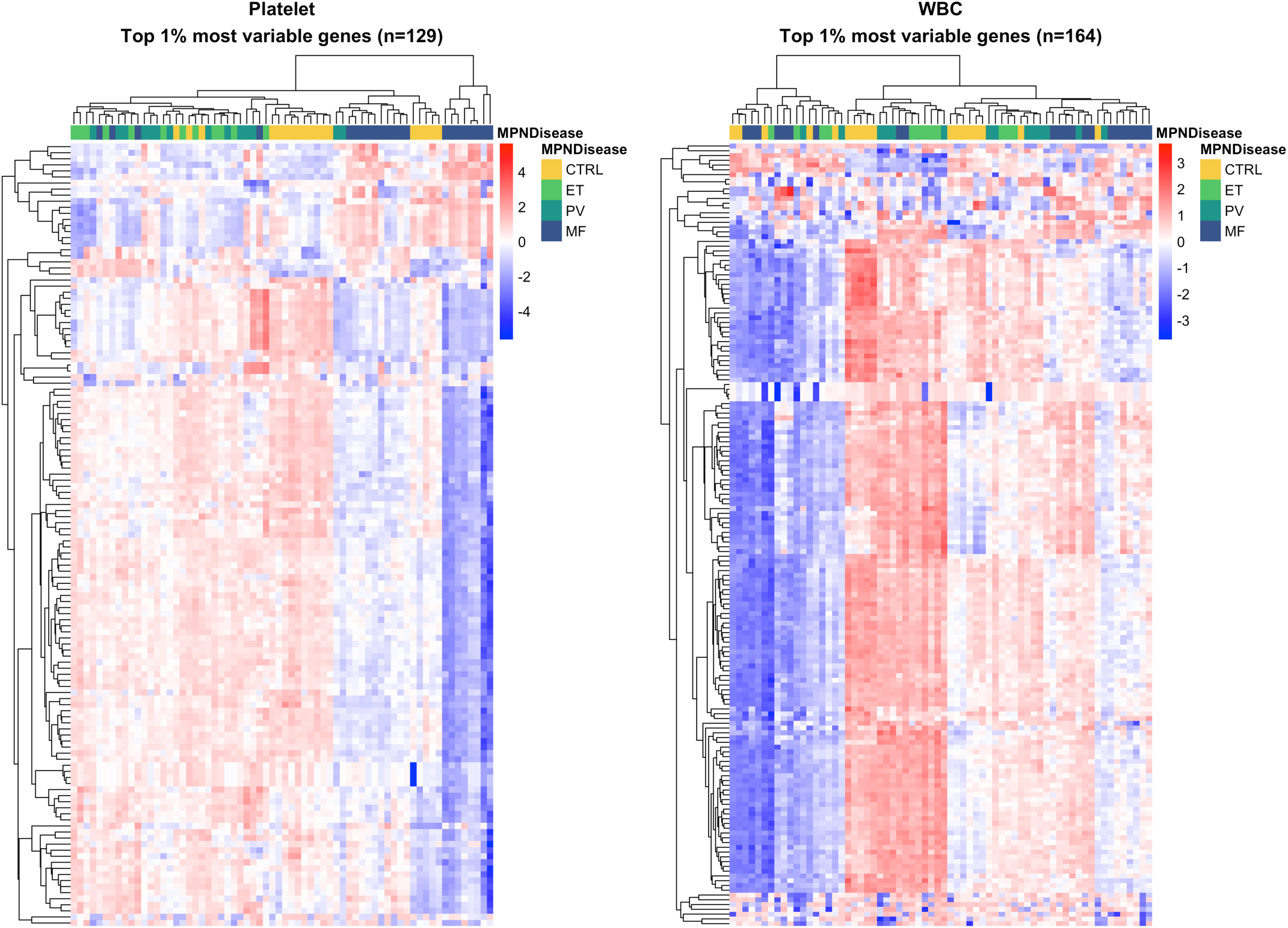
Distinct transcriptional signatures distinguish platelet and WBC responses to myelofibrosis. Side-by-side heatmaps displaying log2-transformed normalized expression values for the top 1% most variable genes (ranked by variance across MF and CTRL samples) in platelet (left panel, n=129 genes) and WBC (right panel, n=164 genes) transcriptomes. Each column represents matched platelet and WBC samples isolated from the same donor blood draw. Rows represent individual genes. Platelet samples were sequenced 2018-2020 (HiSeq 4000); WBC samples were sequenced 2023 (NovaSeq 6000) from cryopreserved aliquots of the identical donor samples. Gene expression is row-scaled (z-score normalization). Color intensity reflects relative expression levels (blue=low, white=intermediate, red=high). Top annotation bar indicates MPN disease subtype (yellow=CTRL, green=ET, teal=PV, dark blue=MF). Hierarchical clustering (Euclidean distance, complete linkage) applied independently to rows and columns reveals clear disease-based sample segregation in both compartments. Platelet samples demonstrate tight clustering with clear separation of all MPN subtypes (MF, PV, ET) from healthy controls. WBC samples show disease vs. control separation but with more heterogeneous expression patterns and greater sample dispersion. Gene sets are non-overlapping between compartments, indicating divergent biological pathways captured by each cell type.

Gene ontology enrichment analysis of platelet-specific differentially expressed genes revealed significant overrepresentation of proteostasis pathways, including endoplasmic reticulum stress response (adjusted p=1.2×10⁻⁸), unfolded protein response (adjusted p=3.4×10⁻⁷), and protein folding machinery (adjusted p=5.1×10⁻⁶). Key genes in these pathways included *CALR*^36,37^*, PDIA6*^38,39^*, HMGA1*^23,40^ - all known components of cellular stress response networks. These findings are consistent with megakaryocyte dysfunction occurring within the fibrotic bone marrow microenvironment, where aberrant cytokine signaling and extracellular matrix remodeling impose sustained proteotoxic stress.

In contrast, WBC-enriched differentially expressed genes highlighted immune activation signatures, including upregulation of Fc gamma receptors (FCGR2A)^41^, death-associated proteins (DAP), and heparanase (HPSE), alongside enrichment of interferon response pathways (adjusted p=2.3×10⁻⁵), cytokine-cytokine receptor interaction (adjusted p=8.7×10⁻⁵), and cell proliferation pathways (adjusted p=1.4×10⁻⁴). These findings suggest that while both compartments reflect MPN pathology, they capture biologically distinct disease dimensions: platelets reflecting proximal bone marrow niche dysfunction, and WBCs reflecting systemic inflammatory responses characteristic of MPN.

### Platelet and WBC Transcriptomes Exhibit Trans-Coregulation Patterns

To investigate whether platelet and WBC transcriptional programs are coordinated across patients, we performed weighted gene co-expression network analysis (WGCNA)^42^ to identify gene modules within each compartment and assess cross-compartment correlations (**Figure 3**). This analysis identified 21 platelet modules and 11 WBC modules. Trans-coregulation analysis revealed 39 significant cross-compartment module correlations (p<0.05, |r|>0.3; Supplementary Table 2). The strongest positive correlation was observed between platelet MEturquoise and WBC MEbrown modules (r=0.52, p=6.8×10⁻⁶), while the strongest negative correlation occurred between platelet MEblue and WBC MEpurple modules (r=-0.52, p=8.3×10⁻⁶). These findings demonstrate that platelet and WBC transcriptional programs exhibit moderate but significant coordination, with both concordant and discordant relationships suggesting complex cross-compartment signaling. The moderate correlation strength (r∼0.5) explains why the combined platelet+WBC classification model (AUROC=0.80) showed intermediate performance between platelet-only (AUROC=0.85) and WBC-only (AUROC=0.77) models, indicating complementary but partially redundant information content.

**FIGURE 3.**
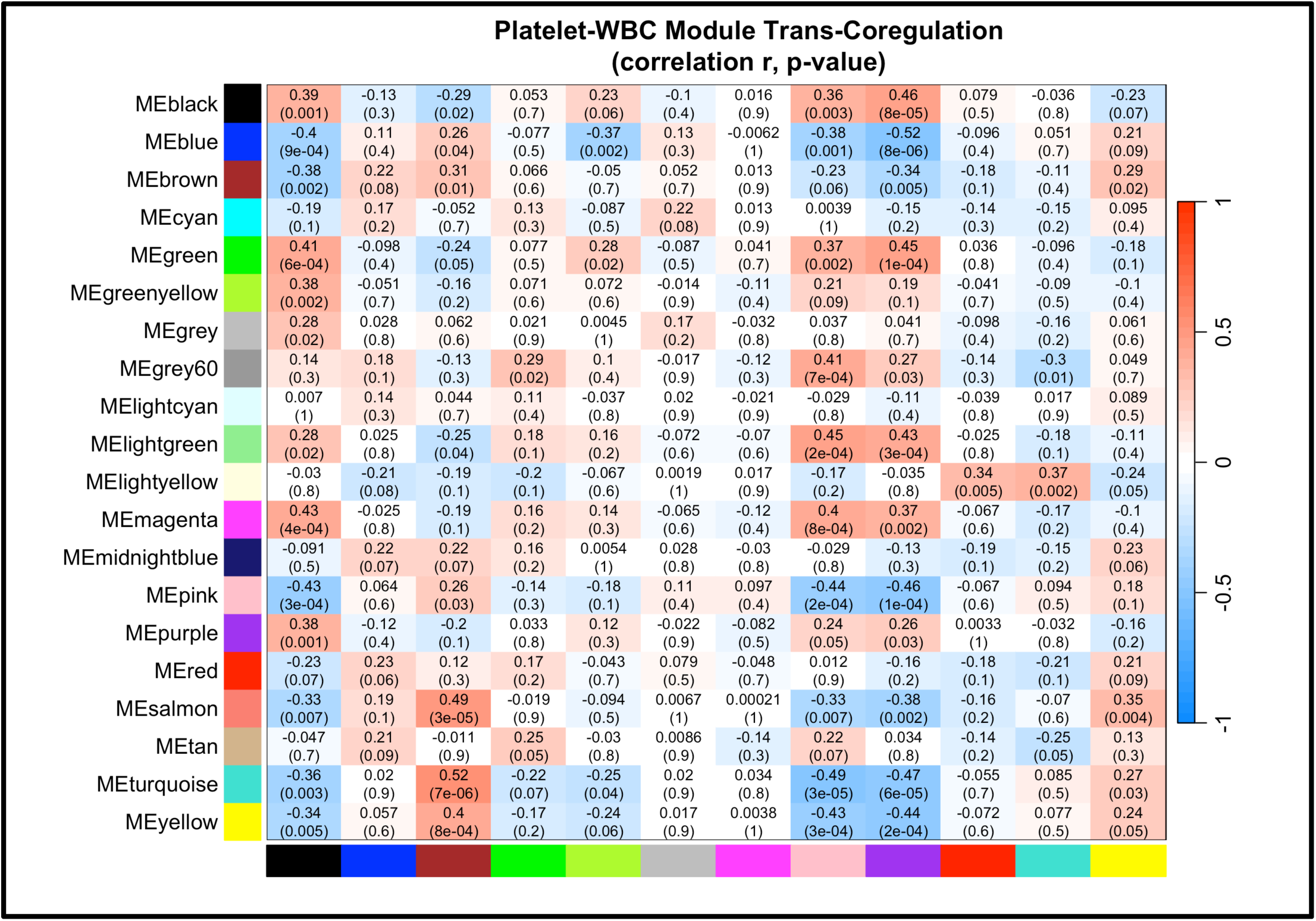
Weighted gene co-expression network analysis reveals coordinated transcriptional programs between platelet and WBC compartments in MPN. Heatmap displaying Pearson correlation coefficients (r) between platelet gene co-expression module eigengenes (rows, 21 modules) and WBC module eigengenes (columns, 11 modules) across 66 matched MPN patient samples. Each cell shows the correlation coefficient (upper value) and corresponding p-value (lower value, in parentheses). Color scale indicates correlation direction and magnitude: red, positive correlation; blue, negative correlation. Highlighted cells indicate significant cross-compartment correlations (p<0.05, |r|>0.3). The strongest positive correlation was observed between platelet MEturquoise and WBC MEbrown (r=0.52, p=7×10⁻⁶), both enriched for proteasome-mediated protein degradation pathways, indicating systemic proteostasis dysfunction across cell compartments. The strongest negative correlation was between platelet MEblue and WBC MEpurple (r=-0.52, p=8×10⁻⁶); platelet MEblue is enriched for cytoplasmic translation and ribosome biogenesis while WBC MEpurple is enriched for mitotic cell division, consistent with but not establishing a compensatory myeloproliferative response. Module colors correspond to WGCNA color labels assigned during network construction. A total of 39 significant cross-compartment correlations were identified (p<0.05, |r|>0.3; Supplementary Table 2).

Pathway enrichment analysis of the most strongly correlated modules revealed coordinated proteostasis dysfunction across compartments. Both platelet MEturquoise and WBC MEbrown—which exhibited strong positive correlation (r=0.52, p=6.8×10⁻⁶)—were enriched for proteasome-mediated protein degradation pathways (adjusted p=0.09 and p=0.01, respectively), indicating systemic proteostasis stress affecting both cell types simultaneously. In contrast, platelet MEblue and WBC MEpurple showed strong negative correlation (r=-0.52, p=8.3×10⁻⁶), with platelet MEblue enriched for cytoplasmic translation and ribosome biogenesis (adjusted p<10⁻⁸⁰) while WBC MEpurple was enriched for mitotic cell division and energy metabolism (adjusted p<0.001). This inverse relationship suggests that impaired platelet protein synthesis capacity correlates with increased WBC proliferation, potentially reflecting compensatory myeloproliferation in response to platelet dysfunction.

### Platelet-Based Transcriptomic Classifiers Outperform WBC and Clinical Models

To assess the relative diagnostic and prognostic utility of platelet versus WBC transcriptomes, we constructed independent multinomial LASSO regression models for MPN subtype classification (**Figure 4**). The models predicted disease class probabilities across four categories (CTRL, ET, PV, MF). To evaluate myelofibrosis detection performance specifically, we generated ROC curves using the predicted MF probability from each multinomial model. This analysis demonstrated superior performance of the platelet-based classifier (AUROC = 0.85, 95% CI: 0.76–0.94) compared to the WBC-based classifier (AUROC = 0.77, 95% CI: 0.66–0.88) in distinguishing MF from other disease states. Both transcriptomic models substantially outperformed a baseline clinical model incorporating only age, sex, and mutation status (AUROC = 0.59, 95% CI: 0.46–0.72), confirming that RNA-based biomarkers capture disease biology beyond traditional clinical parameters (**Figure 4**).

**FIGURE 4.**
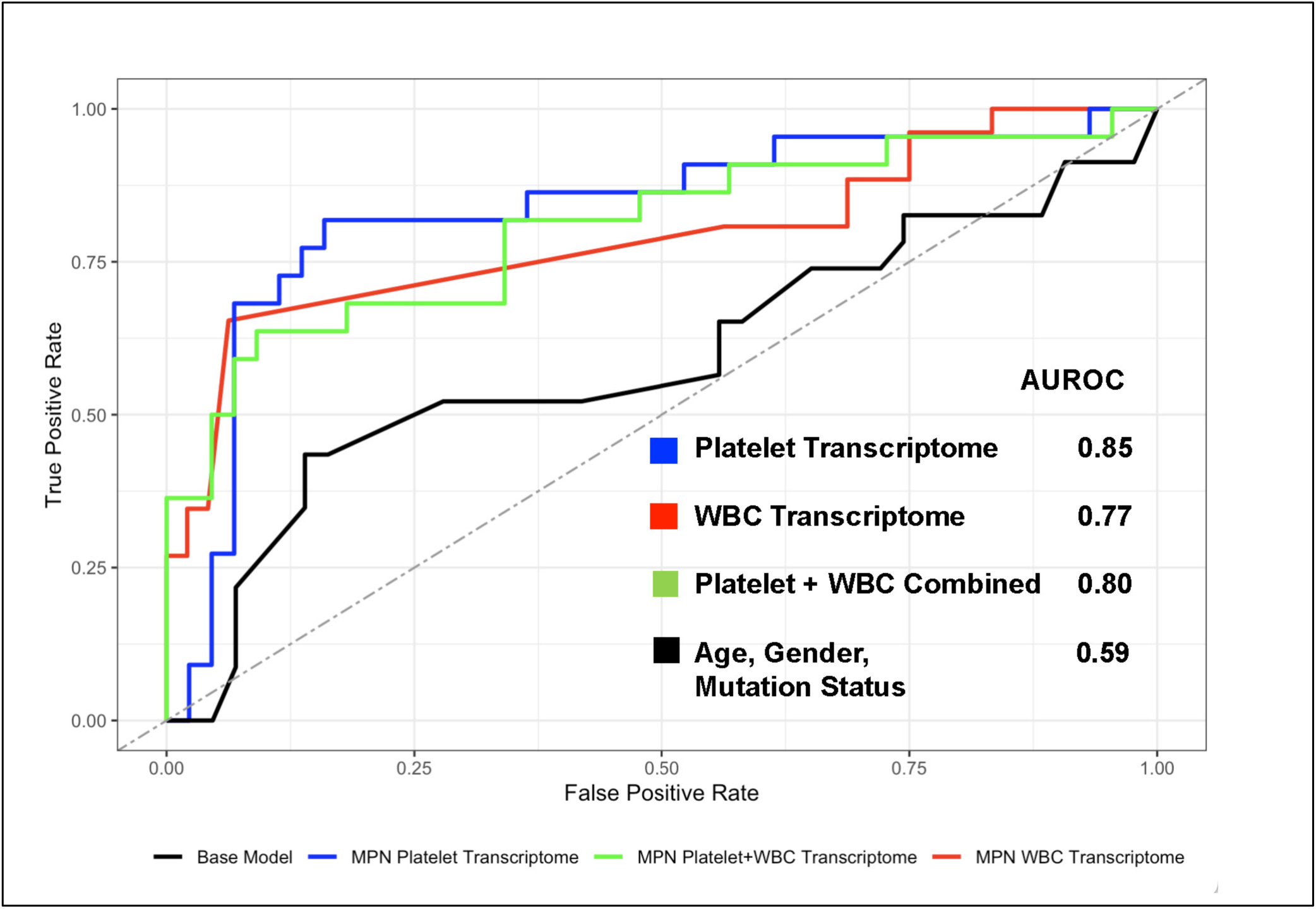
Platelet transcriptomes outperform WBC transcriptomes in myelofibrosis classification. Receiver operating characteristic (ROC) curves comparing four multinomial LASSO regression models for binary classification of myelofibrosis (MF) versus non-MF samples (CTRL, ET, PV combined). Blue curve: platelet transcriptome-based classifier (AUROC=0.85, 95% CI: 0.76-0.94). Red curve: WBC transcriptome-based classifier (AUROC=0.77, 95% CI: 0.66-0.88). Green curve: combined platelet+WBC transcriptome classifier (AUROC=0.80, 95% CI: 0.70-0.90). Black curve: baseline clinical model using age, sex, and mutation status only (AUROC=0.59, 95% CI: 0.46-0.72). Gray dashed diagonal line represents chance performance (AUROC=0.50). The platelet-only model demonstrates superior discriminatory power compared to WBC, clinical, and combined models. Notably, the combined model fails to improve upon platelet-only performance, indicating that platelet features capture the dominant MF-associated signal and WBC features provide complementary but non-additive information. All transcriptomic models substantially outperform clinical parameters alone, confirming that RNA-based biomarkers capture disease biology beyond traditional predictors.

Notably, a combined model integrating both platelet and WBC gene expression features yielded intermediate performance (AUROC = 0.80, 95% CI: 0.70–0.90), failing to exceed the platelet-only model (**Figure 4**). The LASSO models selected entirely distinct gene features from each compartment with no overlap between platelet-selected and WBC-selected panels, consistent with the transcriptome-wide non-overlap described above (88.6% platelet-specific DE genes). That the WBC signal is both biologically distinct and yet non-additive for classification suggests that platelet transcriptomes already capture the dominant MF-discriminating signal, rendering WBC features redundant for this task. This result indicates that platelet and WBC transcriptomes provide complementary but non-additive information.

## DISCUSSION

This study establishes platelet transcriptomic profiling as a superior noninvasive biomarker platform for monitoring myelofibrosis in myeloproliferative neoplasms compared to peripheral blood leukocyte-based approaches. Through comparative analysis of matched CD45-depleted platelet-enriched and CD45+ leukocyte-enriched fractions from 76 MPN patients, we demonstrate that platelet RNA signatures capture bone marrow pathology with greater sensitivity, specificity, and biological fidelity than WBC transcriptomes. Despite WBC samples achieving higher sequencing depth and broader gene detection due to platform differences, platelet transcriptomes exhibited 5.1-fold more differential expression in MF patients (3,453 versus 681 genes at adj p <0.05 and log2FC>1) and superior classification performance (AUROC 0.85 versus 0.77). To rule out that this difference reflects reduced multiple testing burden rather than biology, we downsampled WBC genes to match the platelet baseMean expression distribution and reran DESeq2 on the matched gene set (n=12,797 genes); platelet transcriptomes remained substantially more differentially expressed (2,855 vs 262 DEGs; 22.3% vs 2.0%), confirming the compartment difference reflects genuine biological dysregulation. These findings position platelet liquid biopsy as a powerful tool for disease monitoring, prognostication, and potentially for assessing therapeutic response in clinical trials targeting fibrotic progression.

### Biological Basis for Platelet Superiority

The superior performance of platelet-based biomarkers may reflect fundamental differences in cellular origin and disease exposure between platelets and circulating leukocytes. Platelets are anucleate cytoplasmic fragments derived from megakaryocytes residing within the bone marrow niche, where they experience direct and sustained exposure to the pathological microenvironment characteristic of myelofibrosis - including aberrant extracellular matrix deposition, elevated transforming growth factor-β (TGF-β) signaling, chronic inflammatory cytokine stimulation, and disrupted stromal cell function^17,43,44^. The platelet transcriptome represents a molecular “snapshot” of this local diseased environment, faithfully reflecting megakaryocyte stress responses to fibrotic remodeling^21,45^. In contrast, CD45+ peripheral leukocytes comprise a heterogeneous mixture of neutrophils, monocytes, and lymphocytes that are generated in the marrow but rapidly transit to circulation^46^, potentially experiencing only indirect exposure to fibrosis-associated signals through systemically released inflammatory mediators rather than direct contact with the fibrotic niche itself^47–49^.

The pronounced enrichment of proteostasis pathway dysregulation in platelet transcriptomes—including components of the endoplasmic reticulum stress response, protein folding machinery, and proteasome—provides mechanistic insight into megakaryocyte dysfunction in the fibrotic marrow. These findings are in line with emerging evidence that chronic ER stress and unfolded protein response activation are central to fibrogenic cellular reprogramming in multiple organ systems^50,51^. Megakaryocytes in fibrotic marrow likely experience sustained proteotoxic stress due to aberrant cytokine-driven protein synthesis, accumulation of misfolded proteins, and impaired protein quality control mechanisms. Collectively, these data suggest platelet-based proteostasis signatures may serve as accessible biomarkers for tracking fibrotic burden, warranting further clinical investigation.

Notably, trans-coregulation analysis revealed that this proteostasis dysfunction is not confined to the platelet compartment. The positive correlation between platelet and WBC modules both enriched for proteasome-mediated protein degradation (r=0.52, p=6.8×10⁻⁶) suggests that ER stress and proteostasis dysfunction — previously identified as characteristic of MPN megakaryocytes — extends systemically to circulating leukocytes. This systemic proteostasis stress may contribute to the chronic inflammatory state characteristic of MPNs and could represent a therapeutic vulnerability across cell types. Furthermore, the moderate but significant cross-compartment coordination (r∼0.5) mechanistically explains the intermediate classification performance of the combined platelet+WBC model (AUROC=0.80), as the two compartments provide complementary but partially redundant biological information.

### Comparison to Prior Work and Clinical Context

Our findings complement and extend previous transcriptomic studies of MPN^13,17,33,52–54^. Single-cell RNA sequencing studies have identified megakaryocyte-lineage cells as key drivers of bone marrow fibrosis through aberrant production of pro-fibrotic mediators including platelet-derived growth factor (PDGF), TGF-β, and extracellular matrix proteins^14,15,23,40,55^. RNA sequencing of whole bone marrow biopsies has demonstrated fibrosis-associated gene signatures, but these approaches are invasive and may be confounded by variable cellular composition across samples^33^. Our study demonstrates that peripheral blood platelet RNA captures megakaryocyte-derived signals with sufficient fidelity to enable robust disease classification without requiring bone marrow sampling. This noninvasive approach is particularly valuable for longitudinal monitoring, where repeated bone marrow biopsies may be impractical.

Previous studies examining peripheral blood leukocyte transcriptomes in MPN have primarily focused on mutation-driven clonal expansion and inflammatory signaling rather than fibrosis-specific biomarkers^14,15,22,23,40,55^. Our WBC dataset recapitulates these findings, with immune activation pathways (interferon response, cytokine signaling) dominating the differential expression signature. While these systemic inflammatory signals reflect important aspects of MPN pathophysiology, they lack the specificity and magnitude of platelet-derived fibrosis biomarkers^21^. The failure of the combined platelet+WBC model to improve upon platelet-only classification performance suggests that WBC transcriptomes contribute complementary information about systemic inflammation but do not enhance fibrosis prediction beyond what platelets already provide.

### Methodological Considerations and Limitations

A few technical factors warrant consideration when interpreting these results. Platelet and WBC samples were sequenced on comparative protocols but on different platforms at different times (platelet samples 2018-2020 on HiSeq 4000; WBC samples 2023 on NovaSeq 6000), necessitating independent analysis of each dataset. However, this temporal separation does not invalidate comparative performance assessment, as each dataset was analyzed independently using identical statistical frameworks. The higher gene detection in WBC samples (16,383 versus 12,911 genes) likely reflects the improved sensitivity of the NovaSeq 6000 platform rather than biological differences, yet even with this technical advantage, WBCs exhibited fewer fibrosis-associated transcriptional changes. This paradox strengthens rather than weakens our conclusion that platelet transcriptomes are biologically superior for fibrosis monitoring.

The use of cryopreserved WBC samples raises concerns about RNA degradation affecting gene detection or expression quantification. However, all WBC samples met stringent quality control criteria (RIN ≥7.0), and the robust detection of 16,383 genes argues against widespread degradation artifacts. While both platelet and WBC fractions were isolated fresh from identical blood draws, future studies performing simultaneous sequencing on a single platform would eliminate potential batch effects introduced by temporal separation and sequencer differences.

The CD45+ leukocyte fraction analyzed in this study was collected from platelet-rich plasma after initial centrifugation, raising the question of whether this represents total circulating leukocytes or preferentially captures specific subsets. To address this concern, we compared our WBC differential expression signatures with published peripheral blood leukocyte transcriptomic studies in MPN. Comprehensive analysis of ten independent studies^7,13,33,52–54,56–60^ spanning diverse leukocyte populations (CD34+ cells, granulocytes, monocytes, lymphocytes) consistently identified dysregulation of interferon signaling pathways, cytokine-cytokine receptor interactions, pro-inflammatory gene activation, and JAK-STAT pathway upregulation in MPN peripheral blood. Our WBC dataset recapitulates these established signatures, with significant enrichment of interferon response (adjusted p=2.3×10⁻⁵), cytokine-cytokine receptor interaction (adjusted p=8.7×10⁻⁵), and cell proliferation pathways (adjusted p=1.4×10⁻⁴). This concordance suggests that the CD45+ fraction analyzed captures biologically relevant leukocyte responses characteristic of MPN systemic inflammation rather than artifacts of sample processing.

The CD45+ leukocyte fraction represents a heterogeneous mixture of cell types, and single-cell RNA sequencing approaches could reveal whether specific leukocyte subsets (e.g., monocytes) harbor stronger fibrosis-associated signals that are diluted in bulk analysis. Similarly, platelet preparations, while highly pure (>97%), may contain residual megakaryocyte fragments or platelet-leukocyte aggregates that could influence transcriptomic profiles. These platelet-leukocyte aggregates, which represent ∼2-3% of circulating platelets *in vivo* and are elevated in inflammatory conditions including MPN, reflect physiological cell-cell interactions rather than technical contamination and may contribute biologically relevant transcriptional signals^21^. An additional interpretive consideration is that platelets are anucleate with derivative megakaryocyte mRNA^61^, whereas WBCs are nucleated cells with active transcription. This fundamental difference in RNA biogenesis - static mRNA inheritance versus dynamic transcription - may influence transcriptomic signatures independent of disease biology, though our findings of distinct pathway enrichment patterns (proteostasis in platelets, immune activation in WBCs) suggest that both compartments provide biologically meaningful disease information.

### Clinical Implications and Future Directions

The establishment of platelet RNA as a superior fibrosis biomarker has immediate translational potential for MPN clinical care and therapeutic development. Current fibrosis assessment relies on invasive bone marrow biopsies with semi-quantitative histological grading that suffers from sampling error and inter-observer variability. Platelet RNA-based approaches could enable more frequent, noninvasive monitoring of disease progression, particularly valuable for patients with early-stage ET or PV who may develop secondary myelofibrosis over time. Preliminary analyses suggest that platelet transcriptomes may track fibrosis grade as a continuous variable within MF patients, a finding that warrants dedicated investigation in a study designed specifically for continuous biomarker validation.

Beyond diagnosis and monitoring, platelet transcriptomic biomarkers may prove valuable for assessing therapeutic response in clinical trials. JAK1/2 inhibitors such as ruxolitinib improve symptoms and reduce splenomegaly in MF patients but have limited impact on bone marrow fibrosis. Emerging therapies targeting fibrogenic pathways (e.g., TGF-β inhibitors, anti-fibrotic agents) will require robust biomarkers to demonstrate pharmacodynamic effects on marrow fibrosis without requiring serial bone marrow biopsies. Platelet-based signatures could serve as early pharmacodynamic endpoints, potentially accelerating drug development by enabling rapid assessment of target engagement and biological activity.

Future studies should validate these findings in independent cohorts, ideally with prospective collection of fresh platelet and WBC samples eliminating batch effects and cryopreservation artifacts. Longitudinal analyses tracking individual patients over time would establish whether changes in platelet transcriptomic signatures correlate with clinical progression or therapeutic response. Integration of platelet RNA data with complementary biomarker modalities - including plasma proteomics, circulating microRNAs, and bone marrow imaging - could yield multimodal biomarker panels with enhanced performance. Single-cell RNA sequencing of platelet preparations could identify subpopulations of platelets or residual megakaryocyte fragments with particularly strong fibrosis-associated signatures. Finally, mechanistic studies investigating whether modulation of proteostasis pathways in megakaryocytes (e.g., through pharmacological ER stress inhibitors or proteasome modulators) can attenuate fibrosis progression would establish causality and potentially reveal novel therapeutic targets.

### Conclusion

Comparative transcriptomic profiling of matched platelet and leukocyte fractions establishes platelet RNA as the dominant biomarker compartment for monitoring myelofibrosis in myeloproliferative neoplasms. The enrichment of proteostasis pathway dysregulation in platelet transcriptomes directly reflects megakaryocyte stress within the fibrotic bone marrow niche, providing both mechanistic insight and clinically actionable biomarkers. These findings support the development of platelet liquid biopsy approaches for noninvasive fibrosis monitoring, with potential applications in disease prognostication, therapeutic response assessment, and clinical trial endpoint development.

## METHODS

### Ethical Approval and Patient Cohorts

All MPN peripheral blood samples were obtained under written informed patient consent and were fully anonymized. Study approval was provided by the Stanford University Institutional Review Board (Protocol #18329). Peripheral blood was collected from 76 individuals between 2018 and 2020, including 14 patients with essential thrombocythemia (ET), 17 with polycythemia vera (PV), 27 with primary myelofibrosis (MF), and 18 age-matched healthy controls. MPN diagnoses were established using WHO consensus criteria. Bone marrow fibrosis was graded according to the European Consensus grading system (grades 0-3) based on reticulin staining. Clinical data including age, sex, mutation status (JAK2 V617F, CALR, MPL), and treatment history were obtained through electronic medical record review.

Our cohorts included patients receiving diverse treatment regimens reflecting real-world clinical practice: cytoreductive therapies (hydroxyurea, interferon-alpha), JAK inhibitors (ruxolitinib), anti-thrombotic agents (aspirin), or combinations thereof, alongside treatment-naive patients. This treatment heterogeneity was explicitly adjusted for in all differential expression analyses as a covariate alongside age and sex.

### Blood Collection and Cell Fraction Isolation

Whole blood was collected into acid citrate-dextrose (ACD, 3.2%) sterile tubes and processed to purified platelets within 4 hours of collection using established protocols^62^. Briefly, platelet-rich plasma (PRP) was obtained by centrifugation at 200×g for 20 minutes at room temperature. Prostaglandin E1 was added to PRP to prevent exogenous platelet activation. The PRP was centrifuged at 1000×g for 20 minutes, and the platelet pellet was resuspended in warmed (37°C) PIPES saline glucose buffer.

Leukocyte depletion was performed using CD45 immunomagnetic beads (Miltenyi Biotec) according to the manufacturer’s protocol, yielding a CD45-depleted fraction with >99.9% platelet purity (fewer than 3 leukocytes per 10⁷ platelets as confirmed by hemocytometer counting and post-sequencing analysis of PTPRC/CD45 transcript levels). The CD45-depleted platelet-enriched fraction (1×10⁹ cells) was immediately lysed in TRIzol reagent for RNA extraction. The CD45-positive (CD45+) leukocyte-enriched fraction was resuspended in RNAprotect Cell Reagent (Qiagen) and cryopreserved at -80°C for subsequent analysis.

### RNA Extraction and Quality Control

#### Platelet RNA (2017-2020 cohort and sequencing)

Total RNA was extracted from TRIzol-lysed platelets following DNase I treatment (Qiagen). RNA yield was quantified by NanoDrop 2000 spectrophotometry (Thermo Fisher Scientific) measuring absorbance at 260 nm. RNA purity was assessed by 260/280 nm and 260/230 nm absorbance ratios. RNA integrity was evaluated using the Agilent 2100 Bioanalyzer with the RNA 6000 Nano Chip kit (Agilent Technologies). RNA Integrity Number (RIN) was assigned by the Bioanalyzer Expert 2100 software. Only samples with RIN ≥7.0 were selected for library preparation to control for variable RNA quality. All RNA extractions and library preparations were performed by the same technician to minimize technical variability.

#### WBC RNA (2023 sequencing)

Cryopreserved CD45+ cells were rapidly thawed at 37°C and processed using the RNeasy MinElute Cleanup kit (Qiagen, Cat #74204). RNA quality control was performed using identical criteria (RIN ≥7.0).

### RNA Sequencing Library Preparation

#### Platelet libraries (2018-2020)

Ribosomal RNA-depleted sequencing libraries were prepared using the KAPA Stranded RNA-Seq kit with RiboErase (HMR) (Roche/KAPA Biosystems). cDNA libraries were constructed following the Illumina TrueSeq Stranded mRNA Sample Prep Kit protocol with dual indexing. Library size distribution and quality were assessed on the Agilent 2100 Bioanalyzer, and concentrations were determined by Qubit fluorometry (Thermo Fisher Scientific) for normalization and pooling. Twelve individually indexed samples were pooled per sequencing lane and sequenced on an Illumina HiSeq 4000 platform (Patterned flow cell with HiSeq 4000 SBS v3 chemistry) using 2×75 bp paired-end sequencing with a target depth of 40 million reads per sample. Sequencing was performed at the Stanford Functional Genomics Facility. Output BCL files were converted to FASTQ format and demultiplexed.

#### WBC libraries (2023)

Libraries were prepared using the identical KAPA Stranded RNA-Seq kit with RiboErase (HMR) protocol (KAPA Biosystems, Cat #KK8483). Twelve indexed samples were pooled per lane and sequenced on an Illumina NovaSeq 6000 S4 platform using the NovaSeq 6000 S4 Reagent Kit v1.5 (300 cycles, Cat #20028312). Paired-end sequencing (2×75 bp) was performed with a target depth of 40 million reads per sample at the Stanford Functional Genomics Facility.

### Read Alignment and Gene Quantification

FASTQ files were quality-assessed using FastQC v0.11.9. Adapter sequences and low-quality bases (Phred score <20) were trimmed using Trim Galore v0.6.7. Processed paired-end reads were aligned to the human reference transcriptome (GRCh38/hg38, Ensembl release 104) using RSEM v1.3.3 with Bowtie2 v2.4.2. Gene-level expression was quantified as raw gene counts. Genes with fewer than 10 counts across all samples were filtered prior to normalization. A total of 12,911 genes in the platelet dataset and 16,383 genes in the WBC dataset met expression thresholds and were retained for analysis.

### Differential Expression Analysis

Each cell type dataset (platelet and WBC) was analyzed independently using DESeq2 v1.42.0 in R v4.4.1. Raw count matrices were imported and size factor normalization was performed using DESeq2 default parameters. Differential expression analysis compared each MPN subtype (ET, PV, MF) to healthy controls using the Wald test while adjusting for patient age, sex, and treatment status as covariates. Multiple hypothesis testing correction was applied using the Benjamini-Hochberg method to control false discovery rate (FDR). Genes with adjusted p-value (padj) <0.05 and absolute log2 fold change >1 were classified as significantly differentially expressed. Any duplicate gene entries arising from multiple transcript isoforms were resolved by retaining the instance with the lowest adjusted p-value per unique gene symbol.

Weighted gene co-expression network analysis (WGCNA) was performed independently on platelet and WBC normalized count matrices using the WGCNA R package. Prior to network construction, each dataset was filtered to the top 3,000 most variable genes by variance across the cohorts. Data quality was assessed using the goodSamplesGenes function. Signed co-expression networks were constructed using a soft-thresholding power of β=6 for both compartments. Topological overlap matrices (TOM) were computed from gene-gene adjacency matrices, and hierarchical clustering (average linkage) of the dissimilarity TOM (1-TOM) was used to identify gene modules, with a minimum module size of 30 genes. Module eigengenes (MEs) — the first principal component of each module’s expression matrix — were calculated for both platelet and WBC networks. Trans-coregulation was assessed by computing Pearson correlations between all platelet and WBC module eigengene pairs across the matched samples. Significant cross-compartment correlations were defined as p<0.05 and |r|>0.3. Pathway enrichment analysis of significantly correlated modules was performed using gene ontology biological process terms.

### Multinomial LASSO Classification Models

Disease classification models were constructed independently for platelet and WBC transcriptomes to predict MPN subtype (CTRL, ET, PV, MF). Variance-stabilized expression values from DESeq2 were used as input features. Multinomial LASSO regression with elastic net penalty (alpha=1) was implemented using the glmnet R package v4.1-8. Optimal regularization parameters (lambda) were selected via 10-fold cross-validation by minimizing classification error.

Three models were evaluated: (1) platelet transcriptome-based classifier, (2) WBC transcriptome-based classifier, and (3) combined model incorporating features from both compartments. A baseline clinical model was constructed using multinomial logistic regression with covariates available for all subjects including healthy donors (age, sex, and mutation status: JAK2, CALR, MPL). Mutation status was coded as absent in healthy controls.

Model performance was assessed using receiver operating characteristic (ROC) analysis for binary classification of MF versus non-MF samples. Area under the ROC curve (AUROC) values and 95% confidence intervals were computed using the pROC R package v1.18.5.

### Statistical Analysis and Data Visualization

Heatmaps were generated using the pheatmap R package v1.0.12 with hierarchical clustering (Euclidean distance, complete linkage) applied to log2-transformed normalized expression values. Bar charts comparing differential expression counts across disease subtypes were generated using ggplot2. ROC curves were created using ggplot2 v3.5.1 and pROC v1.18.5. All statistical analyses were performed in R Studio 2024.09.0.375. Statistical significance was defined as FDR<0.05 unless otherwise specified.

## Data Availability

Raw RNA sequencing data will be deposited in NIH GEO databases and made publicly available upon manuscript acceptance.

## Author Contributions

A. Krishnan, J. Rowley, and M. Rondina conceived and designed the study and secured funding. J. Wu coordinated and performed sample acquisition and same-day platelet and leukocyte fraction isolation. V. Natu performed RNA isolation and library preparation for both cell types. A. Krishnan coordinated and oversaw sample sequencing. A. Sawalkar performed WBC RNA-seq bioinformatics pipeline including read alignment and gene quantification. Z. Shen and A. Krishnan performed and interpreted differential expression analysis, LASSO classification modeling, and statistical analyses. Z. Shen and A. Krishnan generated all figures. A. Krishnan, J. Rowley, and M. Rondina interpreted the data. M. Rondina and J. Rowley contributed funding for WBC library preparation reagents and collaborative expertise in platelet biology. A. Krishnan wrote and edited the manuscript. Z. Shen, J. Rowley, and M. Rondina critically reviewed and edited the manuscript. All authors approved the final manuscript.

## Acknowledgements

A.K. sincerely thanks Dr. Jason Gotlib for access to specimens from patients with myeloproliferative neoplasms through the Stanford Cancer Institute and the Stanford University MPN Tissue Bank. Additionally, A.K. thanks Cecelia Perkins and Lenn Fechter for support with coordinating patient specimens and chart information, and Dr. James Zehnder for clinical expertise and guidance. All authors thank the patients at the Stanford Cancer Center for their generous participation in this research. We also thank the cores - Stanford Genomics for RNA sequencing services, technical support and genomic data storage, and additional support from the University of Utah High-Throughput Sequencing Core.

This work would not have been possible without US National Institutes of Health grants 3UL1TR001085-04S1, 1K08HG010061-01A1, 1R21HL181649-01 and the MPN Research Foundation Challenge Grant initiative to A.K, the US National Institutes of Health grants HL155856 and R35HL145237 to M.T.R, and R01HL166805 AND R01HL144957 to J.R..

## Ethics Declarations/Competing Interests

M.T.R. and J.R. hold patent(s) using platelet RNA expression for disease detection.

---

**SUPPLEMENTARY TABLE 1.**
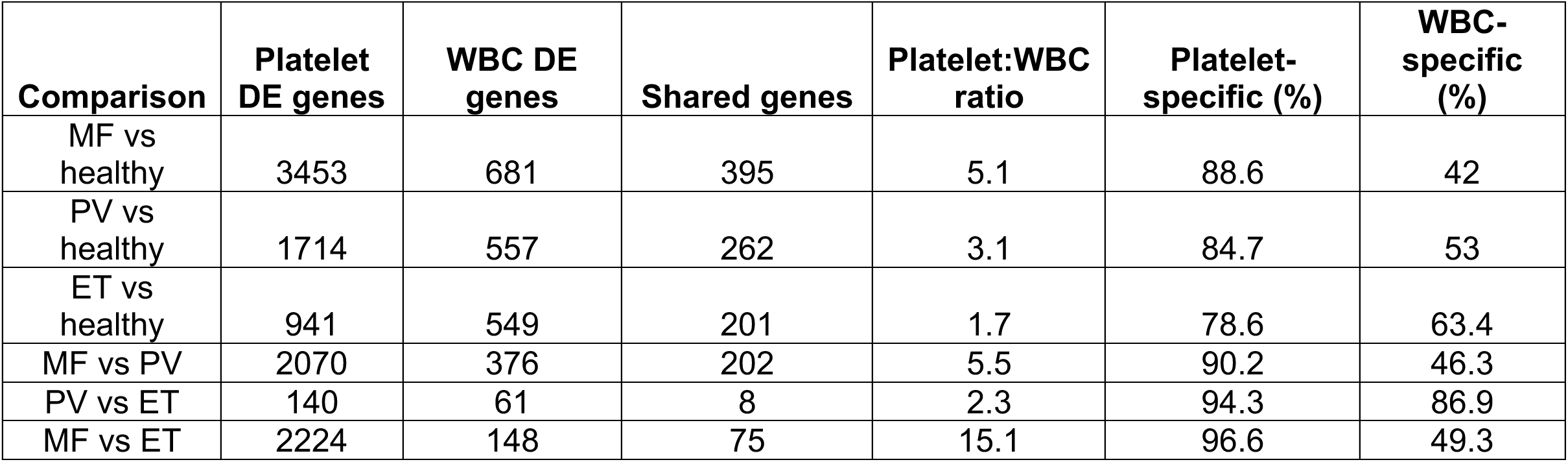

| Comparison | Platelet DE genes | WBC DE genes | Shared genes | Platelet:WBC ratio | Platelet-specific (%) | WBC-specific (%) |
| --- | --- | --- | --- | --- | --- | --- |
| MF vs healthy | 3453 | 681 | 395 | 5.1 | 88.6 | 42 |
| PV vs healthy | 1714 | 557 | 262 | 3.1 | 84.7 | 53 |
| ET vs healthy | 941 | 549 | 201 | 1.7 | 78.6 | 63.4 |
| MF vs PV | 2070 | 376 | 202 | 5.5 | 90.2 | 46.3 |
| PV vs ET | 140 | 61 | 8 | 2.3 | 94.3 | 86.9 |
| MF vs ET | 2224 | 148 | 75 | 15.1 | 96.6 | 49.3 |

**SUPPLEMENTARY TABLE 2.**

| **Platelet_Module** | **WBC_Module** | **Correlation** | **P_value** |
| --- | --- | --- | --- |
| MEturquoise | MEbrown | 0.522398498 | 6.82E-06 |
| MEblue | MEpurple | -0.518157111 | 8.34E-06 |
| MEturquoise | MEpink | -0.490958055 | 2.85E-05 |
| MEsalmon | MEbrown | 0.487831074 | 3.26E-05 |
| MEturquoise | MEpurple | -0.47172953 | 6.38E-05 |
| MEblack | MEpurple | 0.464798636 | 8.43E-05 |
| MEpink | MEpurple | -0.455208941 | 0.000123 |
| MEgreen | MEpurple | 0.453918668 | 0.000129 |
| MElightgreen | MEpink | 0.44516023 | 0.00018 |
| MEyellow | MEpurple | -0.440491253 | 0.000214 |
| MEpink | MEpink | -0.438844473 | 0.000228 |
| MElightgreen | MEpurple | 0.429237797 | 0.000323 |
| MEpink | MEblack | -0.428421447 | 0.000332 |
| MEyellow | MEpink | -0.42783965 | 0.000339 |
| MEmagenta | MEblack | 0.426694651 | 0.000353 |
| MEgreen | MEblack | 0.410809717 | 0.000612 |
| MEgrey60 | MEpink | 0.40628077 | 0.000712 |
| MEyellow | MEbrown | 0.403974069 | 0.000769 |
| MEmagenta | MEpink | 0.403210651 | 0.000788 |
| MEblue | MEblack | -0.399032812 | 0.000904 |
| MEblack | MEblack | 0.394563423 | 0.001044 |
| MEblue | MEpink | -0.38477532 | 0.001423 |
| MEpurple | MEblack | 0.384490983 | 0.001435 |
| MEbrown | MEblack | -0.382483255 | 0.001528 |
| MEgreenyellow | MEblack | 0.382407147 | 0.001531 |
| MEsalmon | MEpurple | -0.379469496 | 0.001676 |
| MEblue | MEgreenyellow | -0.370354952 | 0.002206 |
| MEgreen | MEpink | 0.369951585 | 0.002233 |
| MEmagenta | MEpurple | 0.367149336 | 0.002426 |
| MElightyellow | MEturquoise | 0.366602107 | 0.002465 |
| MEturquoise | MEblack | -0.362098397 | 0.002812 |
| MEblack | MEpink | 0.360276172 | 0.002964 |
| MEsalmon | MEyellow | 0.347839026 | 0.004212 |
| MEbrown | MEpurple | -0.342764689 | 0.004843 |
| MElightyellow | MEred | 0.342389339 | 0.004892 |
| MEyellow | MEblack | -0.340030927 | 0.005216 |
| MEsalmon | MEpink | -0.330196459 | 0.006776 |
| MEsalmon | MEblack | -0.329397077 | 0.006919 |
| MEbrown | MEbrown | 0.312489075 | 0.010634 |

**SUPPLEMENTARY FIGURE 1.**
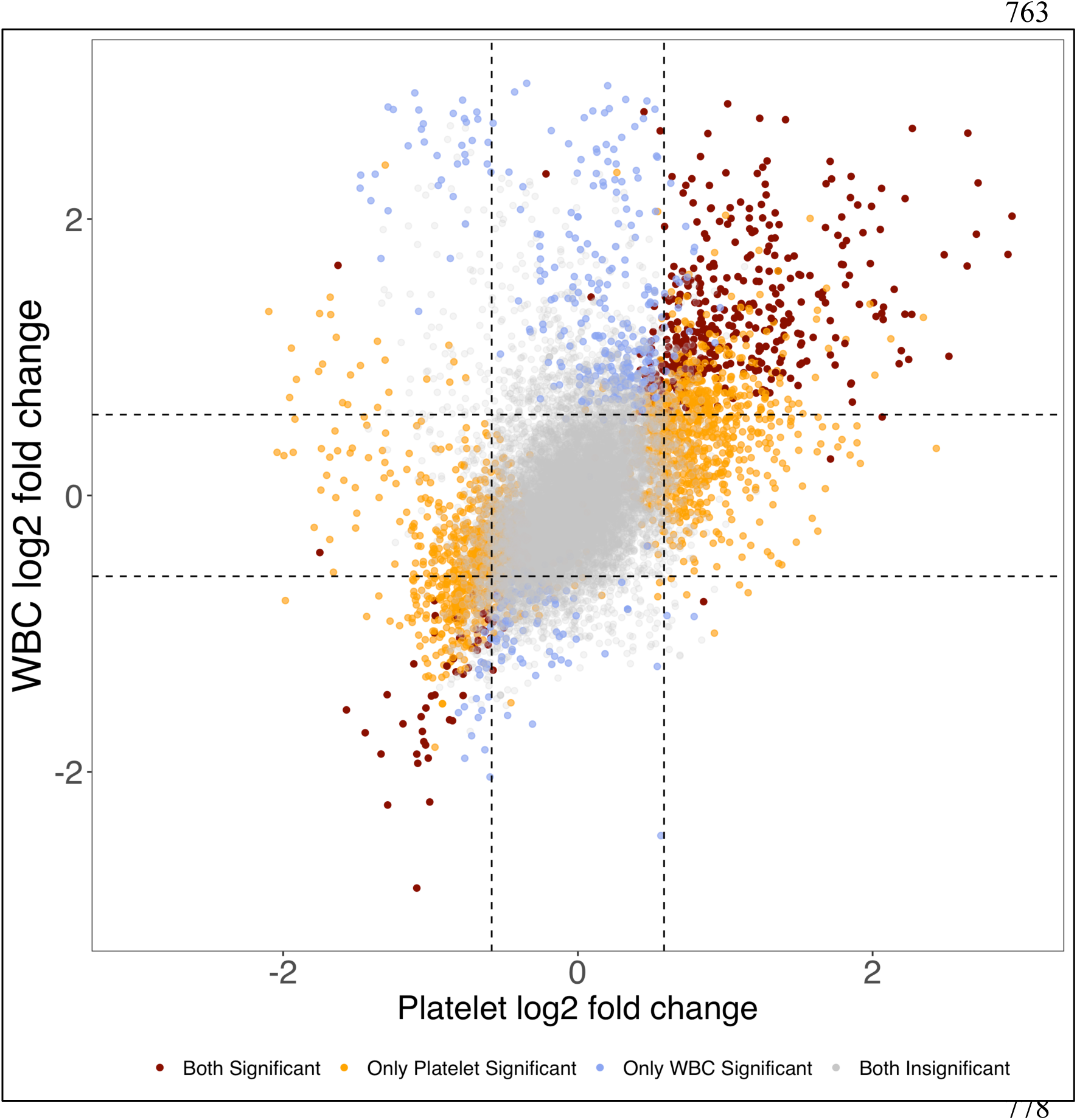
Differential expression concordance between platelet and WBC transcriptomes in myelofibrosis. Scatterplot comparing log2 fold changes (MF vs CTRL) for all detected genes in platelet (x-axis) and WBC (y-axis) datasets. Genes are color-coded by significance: dark red = significant in both compartments (padj<0.05, |log2FC|>1); orange = significant only in platelets; blue = significant only in WBCs; gray = not significant in either. Dashed lines indicate fold change thresholds (±1). Genes significant in both compartments show concordant directionality (clustering along the diagonal), while platelet-specific genes (orange) vastly outnumber WBC-specific genes (blue), consistent with the 5.1-fold difference in differential expression magnitude.

